# Enkurin: A novel marker for myeloproliferative neoplasms from platelet, megakaryocyte, and whole blood specimens

**DOI:** 10.1101/2023.01.07.523111

**Authors:** Sumanth Mosale Seetharam, Yi Liu, Jason Wu, Lenn Fechter, Kanagavel Murugesan, Holden Maecker, Jason Gotlib, James Zehnder, Ramasamy Paulmurugan, Anandi Krishnan

**Author notes:** These authors contributed equally (Sumanth Mosale Seetharam, Yi Liu, Jason Gotlib, James Zehnder).

## Abstract

Impaired protein homeostasis, though well established in age-related disorders, has been linked in recent research with the pathogenesis of myeloproliferative neoplasms (MPNs). As yet, however, little is known about MPN-specific modulators of proteostasis, thus impeding our ability for increased mechanistic understanding and discovery of additional therapeutic targets. Loss of proteostasis, in itself, is traced to dysregulated mechanisms in protein folding and intracellular calcium signaling at the endoplasmic reticulum (ER). Here, using *ex vivo* and *in vitro* systems (including *CD34*^*+*^ cultures from patient bone marrow, and healthy cord/peripheral blood specimens), we extend our prior data from MPN patient platelet RNA sequencing, and discover select proteostasis-associated markers at RNA and/or protein levels in each of platelets, parent megakaryocytes, and whole blood specimens. Importantly, we identify a novel role in MPNs for enkurin (*ENKUR*), a calcium mediator protein, implicated originally only in spermatogenesis. Our data reveal consistent *ENKUR* downregulation at both RNA and protein levels across MPN patient specimens and experimental models, with a concomitant upregulation of a cell cycle marker, *CDC20*. Silencing of *ENKUR* by shRNA in CD34^+^ derived megakaryocytes further confirm this association with *CDC20* at both RNA and protein levels; and indicate a likely role for the *PI3K/Akt* pathway. The inverse association of *ENKUR* and *CDC20* expression was further confirmed upon treatment with thapsigargin (an agent that causes protein misfolding in the ER by selective loss of calcium) in both megakaryocyte and platelet fractions at RNA and protein levels. Together, our work sheds light on enkurin as a novel marker of MPN pathogenesis beyond the genetic alterations; and indicates further mechanistic investigation into a role for dysregulated calcium homeostasis, and ER and protein folding stress in MPN transformation.

**VISUAL ABSTRACT:** 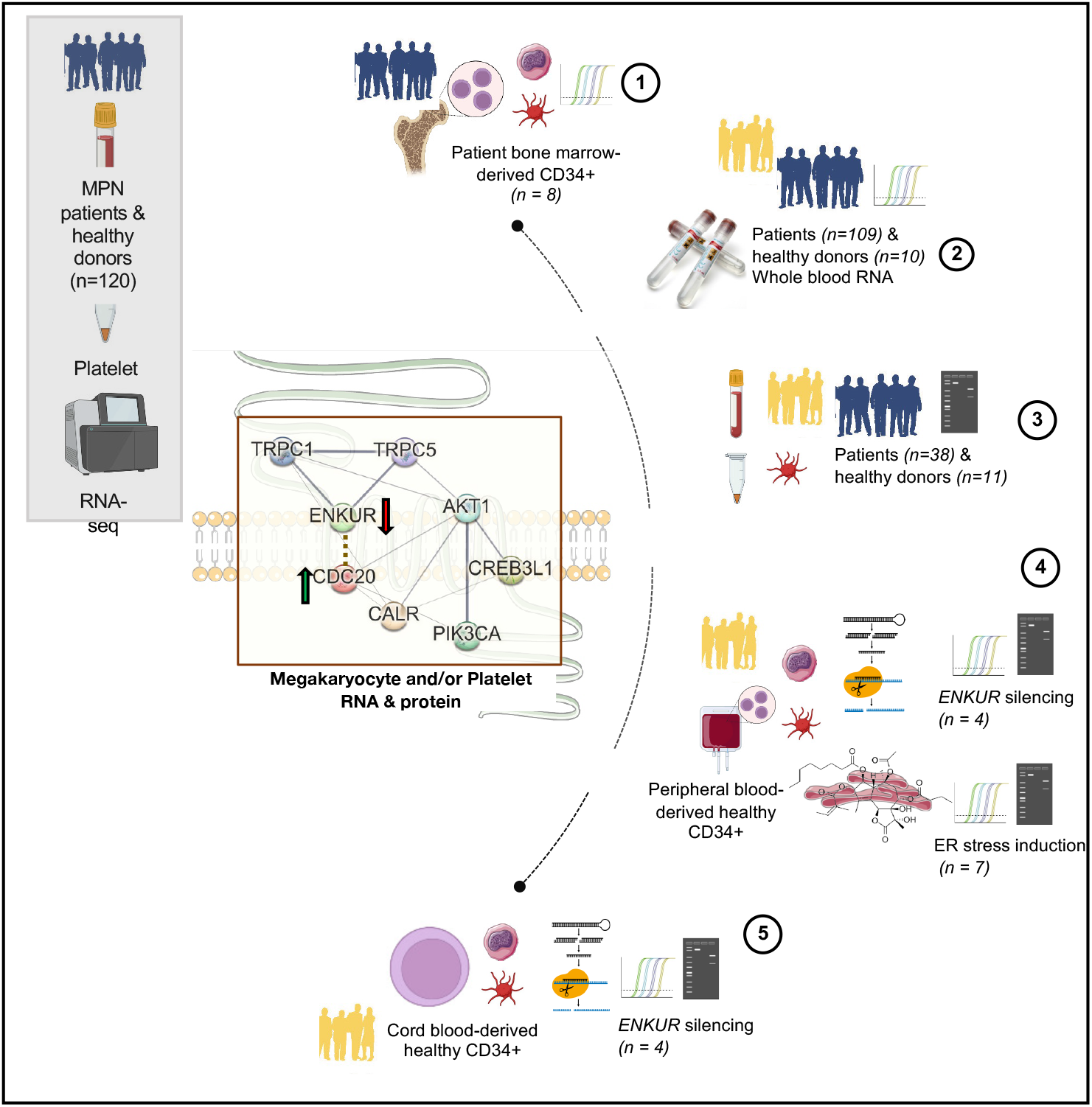

**Key Points:**

1. Enkurin, a calcium adaptor protein, is identified as a novel marker of pathogenesis in MPNs.
2. MPN megakaryocyte and platelet expression of enkurin at RNA and protein levels is inversely associated with a cell differentiation cycle gene, CDC20.
3. Likely role for dysregulated calcium homeostasis, and ER and protein folding stress in MPN transformation.

## Introduction

Myeloproliferative neoplasms (MPNs) are a group of malignant disorders of the bone marrow where a dysregulated balance between proliferation and differentiation gives rise to abnormal numbers of blood cells in circulation. Classical MPNs^1,2^ are defined by a combination of clinical, laboratory, morphological and molecular genetic features, and constitute three phenotypic subtypes: essential thrombocythemia (ET), polycythemia vera (PV), and myelofibrosis (MF, primary or secondary if transformed from prior ET/PV). Somatic mutations in one of three driver genes (*JAK2, CALR, MPL*) constitute their shared molecular genetic pathogenesis; causing constitutive JAK/STAT signaling in hematopoietic stem cells (HSC)^3-7^. The mutated *CALR* also induces oncogenic effects by binding and constitutively activating the thrombopoietin receptor and the downstream signaling cascade^7,8^ causing cellular transformation and abnormal megakaryopoiesis. To date, therapeutic strategies have largely focused on *JAK2* inhibition (e.g. ruxolitinib^9^) and are effective in alleviating MPN symptoms, but only partially, given their primary role in suppression of inflammation and reduction of circulating pro-inflammatory cytokines^10,11^. Additional therapeutic strategies that target MPN pathogenetic mechanisms^12-17^ would be critical in resolving patient disease burden.

A recent study^18,19^ assessing mechanisms of MPN pathogenesis identifies dysregulation of proteostasis and protein quality control at the endoplasmic reticulum (ER) as crucial transformative events, and therefore, a strong therapeutic target. However, this study was primarily in mice and remains to be confirmed in independent additional MPN models and patient-derived specimens. Our recent investigation, profiling the blood platelet transcriptome in all three subtypes of MPN patients^20^ identified high expression of genes and pathways associated with impaired proteostasis, ER stress and unfolded protein response (UPR) well-recognized in other age-related disorders^21-25^. Here, we extend our prior MPN patient platelet transcriptomic data with a focus on genes associated with proteostasis, cell proliferation, ER stress or calcium signaling at high statistical significance (FDR <0.01), and representing four types of trends in platelet RNA expression across MPN subtypes: progressively downregulated, progressively upregulated, consistently upregulated, and uniquely upregulated in MF alone. Genes selected include i) enkurin, *ENKUR*^26^, a little-studied TRPC (transient receptor potential cation channel^27,28^) adaptor protein with known function primarily in the context of sperm motility alone^26,29^, ii) *CALR, a* well-known ER chaperone and calcium binding protein^30-34^, also relevant in the context of megakaryocyte to platelet differentiation^35-38^, iii) cyclic AMP-responsive element-binding protein *CREB3L1*, an ER-Golgi stress transducer and transcription protein involved in the UPR response^39-42^ and lastly iv) *CDC20*, involved in cell division and proliferation^43-45^, and known to be upregulated in other cancers^45^. Using MPN patient bone marrow- and healthy cord- and peripheral blood-derived CD34+ cells differentiated into megakaryocyte and platelet cultures *ex vivo*, we consistently identify *ENKUR* as a novel peripheral and marrow biomarker in MPNs at both RNA and protein levels. Our data also demonstrate a negative correlation of *ENKUR* with *CDC20* expression in the megakaryocyte/platelet cultures, and in independent banked MPN whole blood RNA specimens. Beyond the utility of enkurin as a potential marker of MPN chronic vs advanced subtypes, more research is warranted to elucidate the mechanisms of its downregulation in MPNs, and how that may target cell proliferation pathways.

## Methods

Please see **Table 1** for a tabulated overview of all RNA and protein measurements in this study.

**Table 1:** Summary of all cell types used, their sources and the respective analyses performed.

| Table 1: Experimental Framework and Components |  |  |  |  |  |  |  |
| --- | --- | --- | --- | --- | --- | --- | --- |
| # | Cell type | Source | Megakaryocyte (MK) culture | ENKUR silencing | ER stress induction | RNA expression (MK and/or Platelet) | Protein expression (MK and/or Platelet) |
| 1 | Platelets | Peripheral blood (n = 49) |  |  |  | X<br>*CALR, CREB3L1, CDC20, ENKUR* | X<br>CALR, CREB3L1, CDC20, ENKUR |
| 2 | CD34<sup>+</sup> | Bone marrow (n = 8) | X |  |  | X<br>*CALR, CREB3L1, CDC20, ENKUR* |  |
| 3 | CD34<sup>+</sup> | Cord blood (n = 4) | X | X |  | X<br>*CALR, CREB3L1, CDC20, ENKUR* | X<br>CALR, CREB3L1, CDC20, ENKUR, PI3K and Akt |
| 4 | CD34<sup>+</sup> | Peripheral blood (n = 7) | X | X | X | X<br>*CALR, CREB3L1, CDC20, ENKUR* | X<br>CALR, CREB3L1, CDC20, ENKUR, PI3K and Akt |

### Ethics Statement

All MPN patient and healthy donor samples were obtained under written informed patient consent and were fully anonymized. Study approval was provided by the Stanford University Institutional Review Board. All relevant ethical regulations were followed.

## Materials

We collected MPN patient bone marrow specimens from the Stanford Cancer Center, and whole blood from healthy donors at the Stanford Blood Center. Cell culture supplies including plates, fetal bovine serum, penicillin, streptomycin, phosphate-buffered saline (PBS), and culture medium were procured from GIBCO BRL (Frederick, MD). Antibodies for CALR, CDC20, p-PI3K, p-Akt, b-Actin and anti-rabbit goat IgG were procured from Cell Signaling Technologies (Danvers, MA), for enkurin from Sigma Aldrich (St. Louis, MO) and CREB3L1 from Thermo Scientific (Rockford, IL). Antibodies for Alexa647-tagged CD41, PE-CD42b, PE-CD61, and PE-CD45 were from BioLegend (San Diego, CA). shRNA for ENKUR silencing were synthesized from Protein and Nucleic acid facility at Stanford University (Stanford, CA). The human HEK 293 FT cells and THP1 human macrophage cells were purchased from the American Type Culture Collection (ATCC, Manassas, VA) and cultured following supplier instructions. The CD34, CD45 microbeads, LS and MS columns and magnetic separators were purchased from Miltenyi Biotech (Cambridge, MA). Cytokines IL6, thrombopoietin TPO, and Flt3 ligand were from PeproTech Inc. (Cranbury, NJ); SCF and SFEM II media was from Stem Cell Technologies (Kent, WA). Lenti-X™ GoStix™ Plus was purchased from TaKaRa Bio Inc. (San Jose, CA).

### Peripheral blood platelet isolation

Peripheral blood was collected in acid citrate-dextrose (ACD, 3.2%) sterile yellow-top tubes (Becton, Dickinson and Co.) and was processed within 4 h of collection for all samples. Platelets were isolated by established^46-49^ purification protocols. Briefly, the ACD-tube whole blood was first centrifuged at 200xg for 20min at room temperature (RT). The platelet-rich plasma (PRP) was removed and Prostaglandin E1 was added to the PRP to prevent exogenous platelet activation. The PRP was then centrifuged at 1000xg for 20min at RT. The platelet pellet was re-suspended in warmed (37 deg C) PIPES saline glucose (PSG). Leukocytes were depleted using CD45^+^ magnetic beads (Miltenyi Biotec). Isolated platelets were further resuspended in Trizol or LDS buffer for RNA (PCR) and protein (Western Blot) analyses.

### RNA extraction and quantification from whole blood (PAXgene tubes)

Whole blood from MPN patients and healthy donors was collected into PAXgene tubes (BD Biosciences) containing RBC lysis buffer. RNA was isolated using PAXgene Blood RNA kit following manufacturer instructions (762164, PreAnalytix, Switzerland).

### CD34^+^ cell isolation from MPN bone marrow and healthy peripheral and cord blood cultures

CD34^+^ cells were isolated^50,51^ using microbeads positive selection (Miltenyi Biotec) from source specimens of peripheral blood or cord blood of healthy donors or bone marrow of MPN patients (collected as part of their clinical care with research consent). Briefly, the RBCs were lysed using RBC lysis buffer and the cells pelleted. Cells were suspended in MACS running buffer and incubated with the CD34^+^ microbeads for 1 hour on ice and then passed through MACS LS columns pre-equilibrated with MACS running buffer. CD34^+^ cells thus collected were washed using MACS running buffer and resuspended in the SFEM II media (4-5×10^5^ cells/mL) containing TPO (20ng/ml), SCF (25ng/ml) and Gentamicin (1:1000) and plated/transferred to 12 well plates at 37 °C in 5% CO_2_. On day 3, cells were resuspended in fresh media (SFEM II) containing TPO (20ng/ml), SCF (25ng/ml) and Gentamicin (1:1000), and on day 6, 9, and 12, in TPO (40ng/ml) and Gentamicin (1:1000). The cells were harvested on day 15. Cells were collected and centrifuged at 300xg for 10 min to pellet the larger megakaryocyte fraction and at 3000xg for 30 min for the smaller platelet fraction. With both peripheral- and cord-blood-derived CD34^+^ cells, the isolated cells were expanded using hematopoietic stem cell expansion media (Cell Genix) containing 100ng/ml of IL6, Flt3 ligand, SCF, and TPO for 5 days before starting the culture. The formation of MK cells and platelets were confirmed by surface marker analysis with CD41, CD42b and CD61.

### Cell surface marker analysis using flow cytometry

Following isolation of the megakaryocyte and platelet fractions from the 14-day (+ TPO) CD34^+^ culture, cells were collected and washed with PBS. For the CD41, CD42b and CD61 stains, cells were resuspended in MACS running buffer and stained with Alexa 647 anti-human CD41, PE anti-human CD42b, and PE anti-human CD61, and incubated at RT protected from light for 45 min. Cells were washed with PBS and resuspended at a final concentration of 2 × 10^7^ cells/ml in PBS, prior to running the sample on the flow cytometer. All experiments were performed using GUAVA Flow Cytometer and all flow cytometry analysis was performed in the FlowJo Software. Data from flow cytometry experiments was acquired by gating for events that were in focus.

### Total RNA isolation and PCR with CD34^+^ derived megakaryocyte and platelet fractions

Total RNA was isolated from the megakaryocyte (MK) and platelet fractions using a mirVana RNA extraction kit (Life Technologies, Grand Island, NY) in accordance with the manufacturer’s instructions. Briefly, MKs/platelets were homogenized into 300μl lysis buffer, then incubated with 30μl homogenate additive for 10 min. RNA was extracted with acid-phenol and column purified, then washed three times with washing buffer, and eluted in 40μl of sterile elution buffer. The total RNA was quantified first using a Nanodrop spectrophotometer, then 100ng of total RNAs were reverse-transcribed using a Reverse-Transcription Kit (Life Technologies) and RT primers to synthesize cDNA. Then, qRT-PCR was performed using the TaqMan-PCR primers and probe mix combined with the cDNA derivates. qRT-PCR was performed through 2 min incubation at 50 °C, then followed by DNA polymerase activation at 95 °C for 10 min; plus 60-cycles at 95 °C for 15 s, and 60 °C for 60 s in the BioRad CFX96 thermocycler system (BioRad). The qRT-PCR reaction procedure was executed in a 20μl final reaction volume. The expression of target genes was analyzed by the 2−ΔΔCt method.

### Thapsigargin-induced ER stress

Isolated human CD34^+^ cells from the peripheral blood of healthy volunteers were cultured for 15 days with the supplementation of both SCF and TPO for the first 6 days and then with TPO alone till day 15. Fresh media containing either SCF and/ or TPO was added every 3 days. To induce ER stress in these cells, known ER stressor drug Thapsigargin (125nM, as optimum dosage identified via preliminary experiments) was added to the culture on day 7. The culture was maintained till day 15 with media change every 3 days. The cells were harvested and analyzed for gene expression and immunoblotting for the target proteins.

### Western Blotting for protein expression in megakaryocytes and platelets

Isolated MKs were aliquoted for flow cytometry, RNA expression and western blotting. For western blotting, an aliquot of the MK cells was pelleted and lysed using cell lysis buffer containing protease inhibitor cocktail, EDTA. The samples were then resolved on a reducing SDS-PAGE (10% acrylamide). Blots were stained using appropriate primary (anti-human rabbit antibody; 1:1000 v/v) and secondary antibodies (anti-rabbit IgG HRP conjugate; 1:2000 v/v).

Protein expression was determined based on the detection of a band. The intensity of the protein bands observed was semi-quantified using IVIS or ImageJ software with normalization of each protein against beta-actin. Blots were visualized using ECL reagent by Amersham Imager 680 or by IVIS imaging systems.

### ENKUR gene silencing

#### a. Construction of the vector

PLKO1 vector was used to construct the ENKUR-silenced cell lines in HEK293T cells (known for their ease of transfection and fast growth rate). The PLKO1 vector was cut using EcoR1 and Age1 to ligate the shRNA using T4 ligase.

The shRNA sequences used were:

shRNA1: PCCGGTCCGGCCAACCTCGATACTCTTATTTCTCGAGAAATAAGAGTATCG AGGTTGGTTTTTTTTGG

shRNA 2: PCCGGTCCGGCATGGGAGTGGCTAAAAAGCCCTCGAGGGCTTTTTAGCC ACTCCCATGTTTTTTTTGG

Scrambled: PCCGGTCCGGGTGCGTTGCTAGTACCAACTCTCGAGAGTTGGTACTA GCAACGCACTTTTTTTTGG

#### b. Validation of the construct

The silencing effect of the shRNA construct was assessed using HEK293 FT cells by transfection of the shRNA containing plasmids (pLKO1-shRNA) using lipofectamine method. Briefly, 100K HEK293 FT cells were seeded in a 12-well plate 24h prior to the transfection. Transfection was performed by adding a master mix containing lipofectamine and the shRNA in reduced serum media for 4 hours, before adding the complete media. The cells were harvested 48h after transfection and the silencing effect assessed by western immunoblotting against enkurin.

### Cell culture for ENKUR silencing

HEK293 FT cells were cultured in Dulbecco’s Modified Eagle’s Medium/high glucose with 10% FBS, 0.1% streptomycin, and 100 U mL^−1^ penicillin at 37 °C, 5% CO_2_, and 95% air environment, while THP-1 cells were cultured in RPMI media containing 10% FBS, 0.1% streptomycin, and 100 U mL−1 penicillin at 37 °C, 5% CO_2_, and 95% air environment. The cells were tested for any mycoplasma contamination using a MycoAlert kit (Lonza, Allendale, NJ), and were maintained at optimum cultural conditions.

### Production of lentiviral particles

After the confirmation of the silencing effect of the shRNA construct, they were packaged along with the viral plasmids to produce the lentivirus. Briefly, the lentiviral vectors were transfected to HEK293 FT cells using calcium phosphate method. The cells were supplemented with chloroquine and HEPES. After 48h of incubation, the media was carefully aspirated, and the viral particles were concentrated by ultracentrifugation. The IFU value was determined by Lenti-X GoStix following manufacturer instructions (Cat#631280).

### Validation of the lentivirus silencing in HEK cells

As above, 100K HEK293 FT cells were plated in a 12-well plate 24h prior to lentiviral infection (transduction). The media was aspirated and lentivirus master mix containing polybrene in reduced serum media was added to the cells. After 4h of incubation, the cells were supplemented with complete media and was incubated for 48h before harvesting. The cells were lysed and immunoblotted against enkurin to assess the gene silencing effect.

### ENKUR gene silencing in CD34^+^ derived megakaryocytes

The shRNA expressing lentiviral particles were infected into the CD34^+^ cells in culture on day 9 (MKs derived from CD34^+^ cells) at a ratio of 1:20 (cell: viral particle). The culture was maintained for another 6 days with supplementation of TPO every 3 days. Cells were harvested on day 15; and centrifuged at two speeds; low (300xg) and high (3000xg) to separate the MK and platelet fractions respectively. Each fraction was divided into 3 aliquots for assessing each of cell surface markers (50 – 100K cells), RNA (∼50K cells) and protein (100 – 200K cells) expression.

### Statistical Analyses

Continuous variables from all experiments were assessed for normality. Data that were normally distributed were expressed as a mean plus or minus the standard error of the mean. For analyses involving two groups, a parametric two-tailed student t-test was used. When three or more groups were analyzed, an ANOVA with a Tukey’s post-hoc test was performed. When data were not normally distributed a Mann-Whitney was used when two groups were analyzed while a Kruskal-Wallis with a Dunn’s multiple comparison post-hoc test was used for analyses of three or more groups. When appropriate a two-way ANOVA with post-hoc test was used as described. Summary statistics were used to describe the study cohort and clinical variables were expressed as the mean ± error of the mean or as a number and percentage (%). Statistical analyses were performed by using GraphPad Prism (version 9, San Diego, CA), and a p-value < 0.05 was considered statistically significant.

## Results

### Validation of RNA and protein expression of 4 MPN candidate markers

Four candidate markers from our prior platelet RNA-seq analyses^20^ were identified based on specific trends in progressive expression across the 3 MPN subtypes vis-à-vis healthy donors (FDR<0.01, n=120) independent of patient driver mutational status. *ENKUR and CALR*, in particular, were identified based on opposing trends in progressive expression (**Figure 1A and Figure S1A**), *CDC20* on its specific high expression in MF patients alone in contrast with healthy donors and ET or PV patients (**Figure 1A**, top right), and *CREB3L1* based on its substantially increased (> 50-fold) expression in all MPNs vs healthy (**Figure S1A**, top right). Given that the platelet RNA profile is a composite of influences from the peripheral circulation as well as of the genetic (and MPN mutational) profile of megakaryocytes in the bone marrow, we hypothesized *a priori* that our MPN candidate markers are likely to be variably validated in one versus the other.

**Figure 1:**
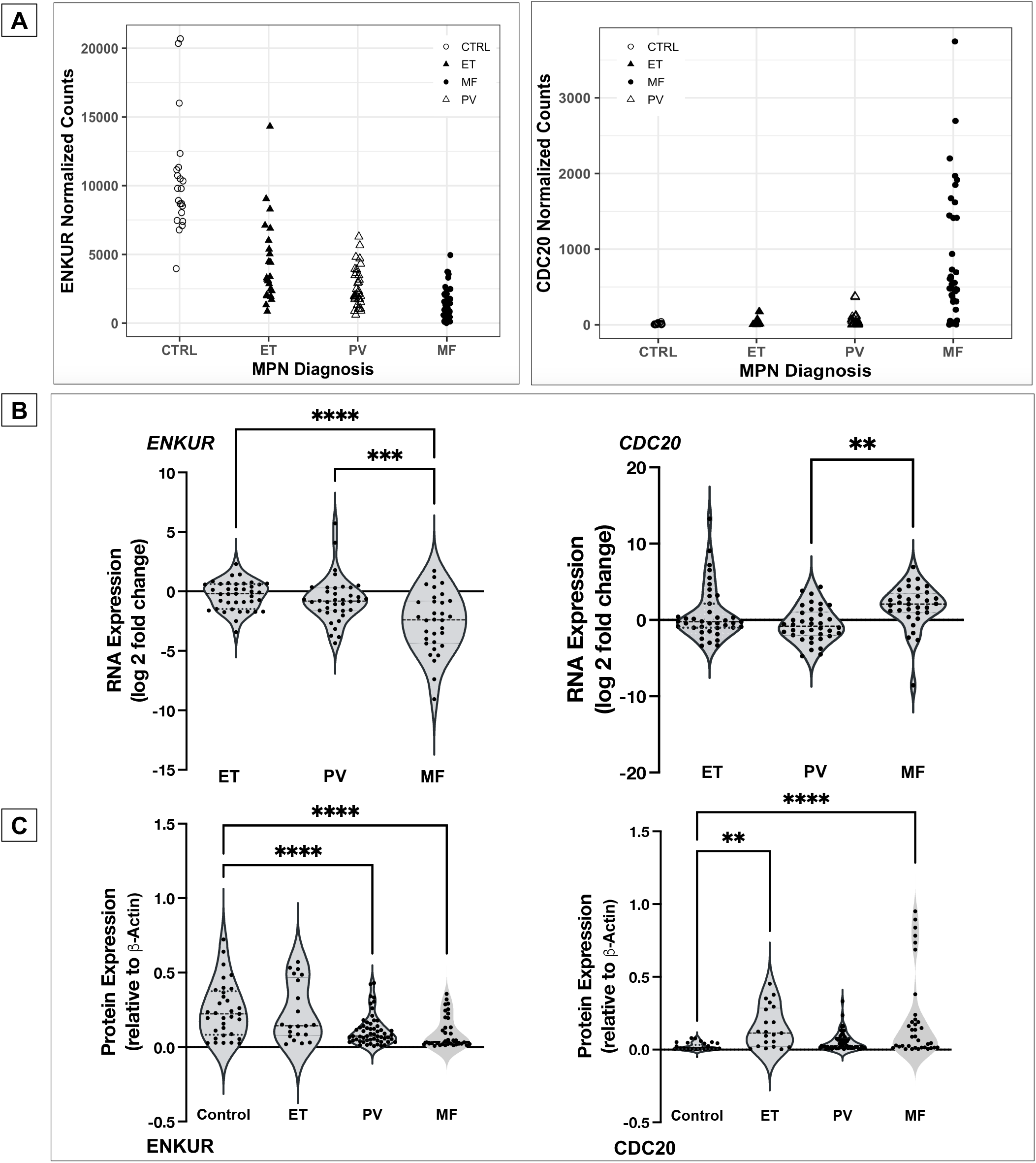
RNA expression in platelets and whole blood of MPN patients and protein expression in platelets. (A) MPN Platelet RNA-seq normalized expression data: MPN patients (total n=99: ET n=24; PV n=33; MF n=42) vs healthy donors (n=21) (B) whole blood RNA: MPN patients (total n=109: ET n=39; PV n=38; MF n=32) vs healthy donors (n=10) (C) platelet protein: peripheral blood platelets were isolated from MPN patients (total n=38: MF n=12; PV n=19; ET n=7) and healthy donors (n=11). Bonferroni multiple comparisons adjusted p values ****p<0.0001, ***p<0.001 and **p<0.01 when compared with controls (healthy donors). All RNA expression is normalized to GAPDH and expressed as log2 fold change.

First, in whole blood RNA of MPN patients versus healthy donors, expression of *ENKUR* and *CDC20* was confirmed by qPCR (**Figure 1B**, n=109, p < 0.005) as nearly 6-fold downregulated and 3-fold upregulated respectively in the advanced MPN subtype, MF. Expression of *CREB3L1* was also confirmed as over 15-fold upregulated across ET, PV, and MF (**Figure S1B**, right panel), whereas expression of *CALR* (**Figure S1B**, left panel) was variable and not uniformly consistent with the platelet transcriptome (likely owing to lower resolution for detection of transcripts in whole blood). Next, in assessing platelet protein levels, trends in expression of ENKUR was significantly different in PV and MF versus healthy donors, and CDC20 in ET and MF (**Figure 1C**), whereas CALR and CREB3L1 were not significantly different between patient and healthy donor specimens (**Figure S1C**).

### Megakaryocyte and platelet fractions from CD34^+^ cultures of MPN patient bone marrow

To further delineate RNA expression of the candidate markers as specific to platelets in circulation alone or also in marrow-derived megakaryocytes, we established an *ex vivo* culture of CD34^+^ stem cells from MPN patient bone marrow (n=8, 4 MF) and generated megakaryocyte and platelet fractions (**Figure 2A-B**, and Methods and **Figure S2A-B** for flow cytometric and histological confirmation of cultured megakaryocyte and platelet fractions). Given the variable differentiation states of the nascent platelet fraction and its lower cell quantity than the megakaryocytic fraction, we focused entirely on the megakaryocytes harvested. Downregulation of *ENKUR* was recapitulated in these patient-derived megakaryocytic RNA (mean approx. 2-fold reduction in MF alone and 5-fold all MPN) with associated upregulation of *CDC20*, CREB3L1 (mean of approx. 1-fold) and *CALR* (∼4-fold) in all MPN (n=8, **Figure 2C**).

**Figure 2:**
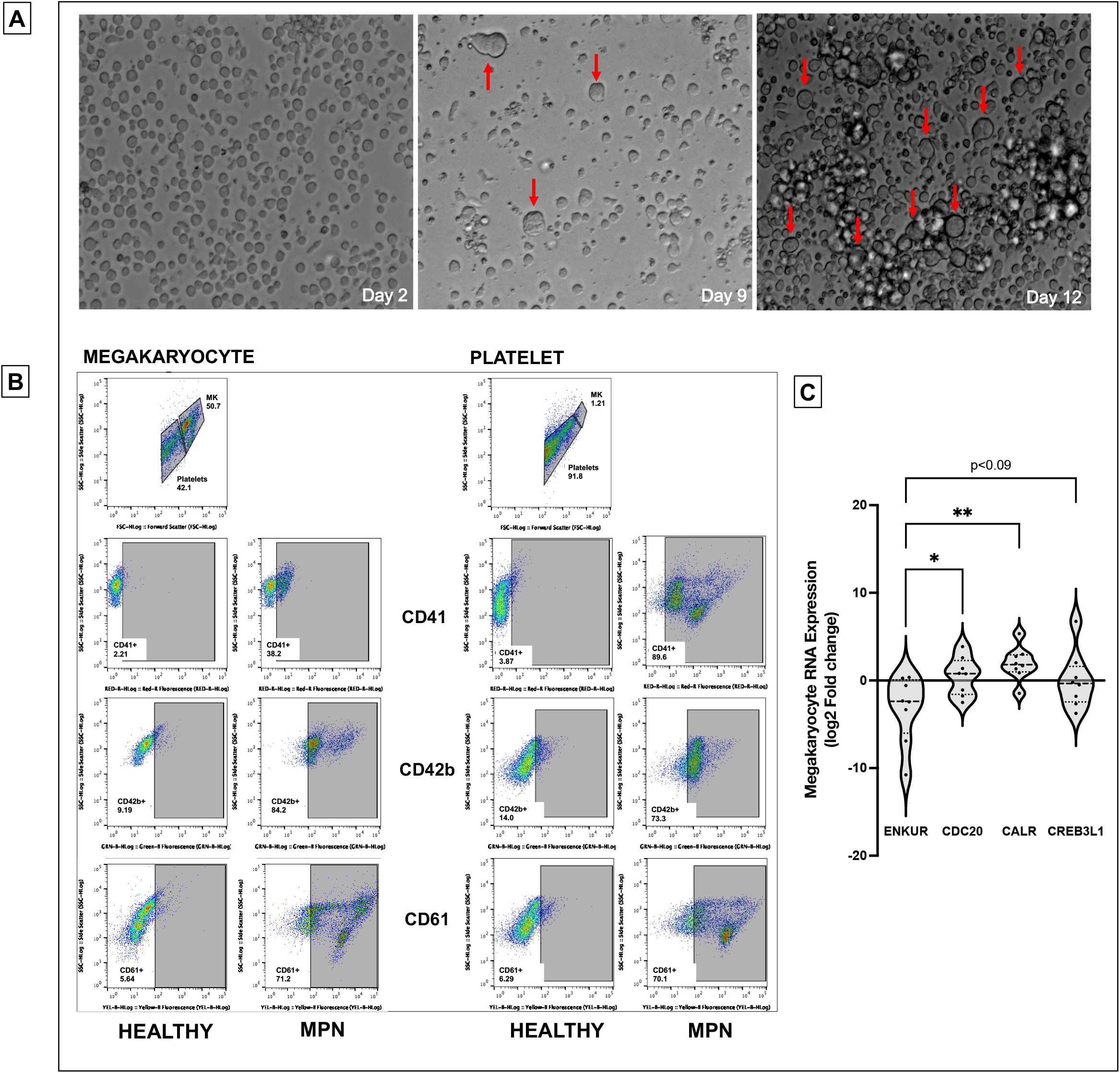
Culturing of CD34^+^ cells from the bone marrow of MPN patients and RNA expression. (A) Microscopic images of the CD34^+^ cells isolated from the bone marrow of MPN patients over the 15-day culture period at 20X magnification. Notice the formation of megakaryocytes indicated by the red arrow. (B) Flow cytometric analysis of cell surface markers to confirm the production of megakaryocyte and platelet fractions after 15 days of culture. (C) megakaryocytic RNA: cell culture from fresh MPN patient bone marrow (n=8) derived CD34^+^ cells then differentiated over 15 days into megakaryocytes. All RNA expression is normalized to GAPDH and expressed as log2 fold change. Bonferroni adjusted p values **p<0.01, *p<0.05.

### *ENKUR* gene silencing

To better evaluate the negative correlation between *ENKUR* and *CDC20/CALR* expression, we sought lentiviral knockdown in primary CD34^+^ cells (derived from healthy peripheral blood and cord blood specimens) using two distinct *ENKUR* shRNA constructs. **Figure 3A-B** describes our experimental framework. The shRNAs were first expressed in pLKO.1 plasmid that was amplified and isolated from *E*.*coli*, and the lentiviral plasmids were then transfected into HEK293 FT cells to generate the lentivirus. *ENKUR* shRNA silencing and downregulation of enkurin protein was confirmed at a concentration of 1:20 HEK cell:viral particle. Primary CD34^+^ cells from healthy peripheral (n=4) and cord blood (n=4) specimens were cultured at four conditions: i) control with no ENKUR silencing, ii) *and* iii) lentiviral knockdown with the two shRNA constructs, and finally iv) scramble control shRNA. Where relevant, shRNA infection was introduced at day 7 of culture and continued for another 7 days (totaling a 15-day culture, **Figure 3B**). Megakaryocyte and platelet fractions thus derived were confirmed by flow cytometric analysis (**Figure 4A**). Downregulated RNA expression of *ENKUR* was significantly associated with concomitant high expression of *CDC20* in both of the megakaryocyte and platelet fractions (**Figure 4B**). Cell quantity in the platelet fraction was just sufficient for RNA qPCR measurements alone, and therefore, protein levels could be assessed only in the megakaryocytes. Inverse correlation between ENKUR and CDC20 was validated even in megakaryocytic protein expression (**Figure 4C)**. However, *ENKUR* silencing alone (without the associated patient factors in our *in vitro* experiments, e.g. mutational status) was insufficient to generate a statistically significant differential in *CALR or CREB3L1* expression at both RNA and protein levels (**Figure S3A-B**).

**Figure 3:**
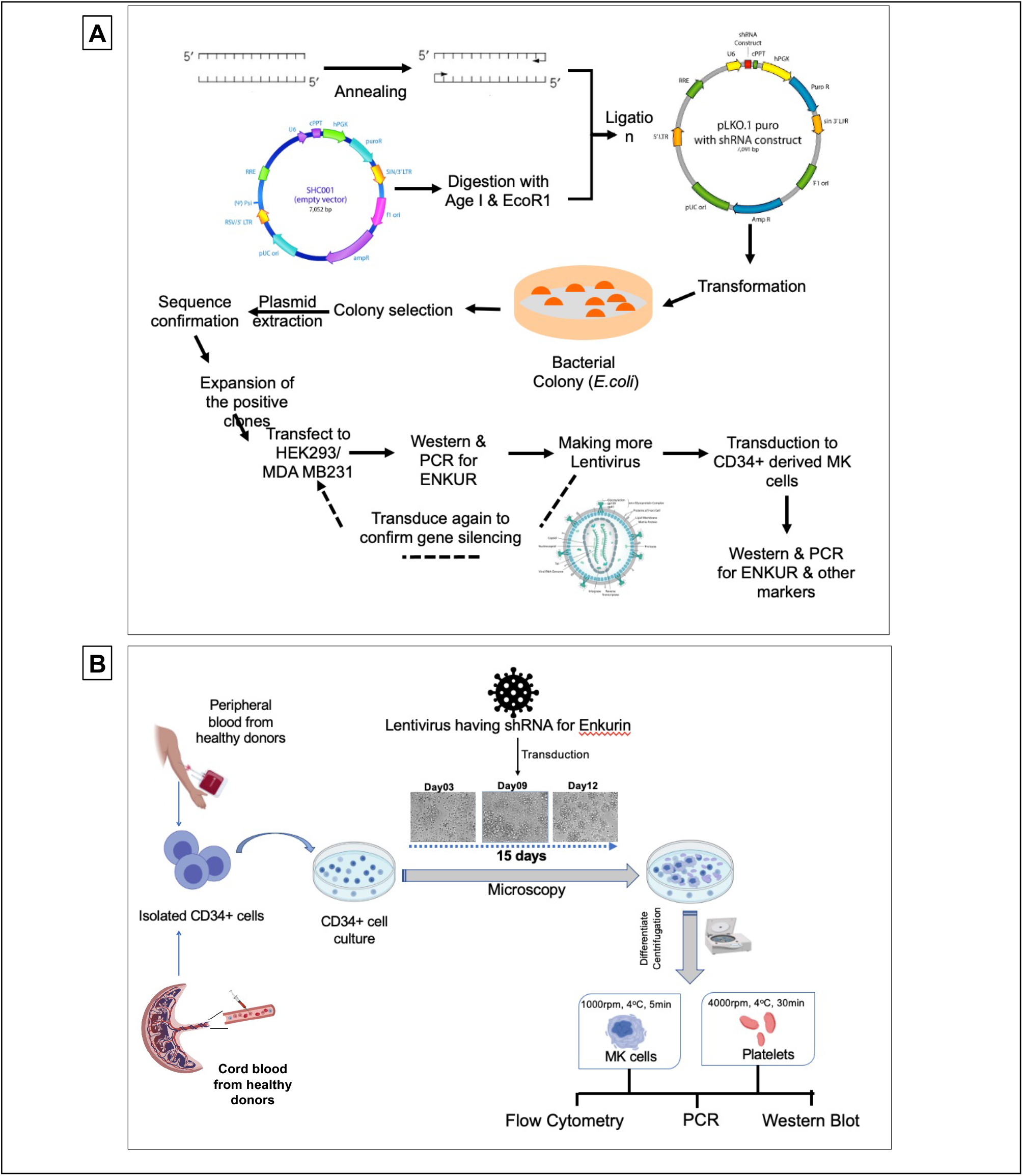
Experimental design and workflow of Lentiviral transduction-induced silencing of ENKUR gene in CD34^+^ stem cells. (A) Workflow of the shRNA construction and generation of the lentivirus (B) experimental plan for the silencing of the ENKUR gene in CD34^+^ stem cells. The CD34^+^ cells were isolated from both peripheral and cord blood.

**Figure 4:**
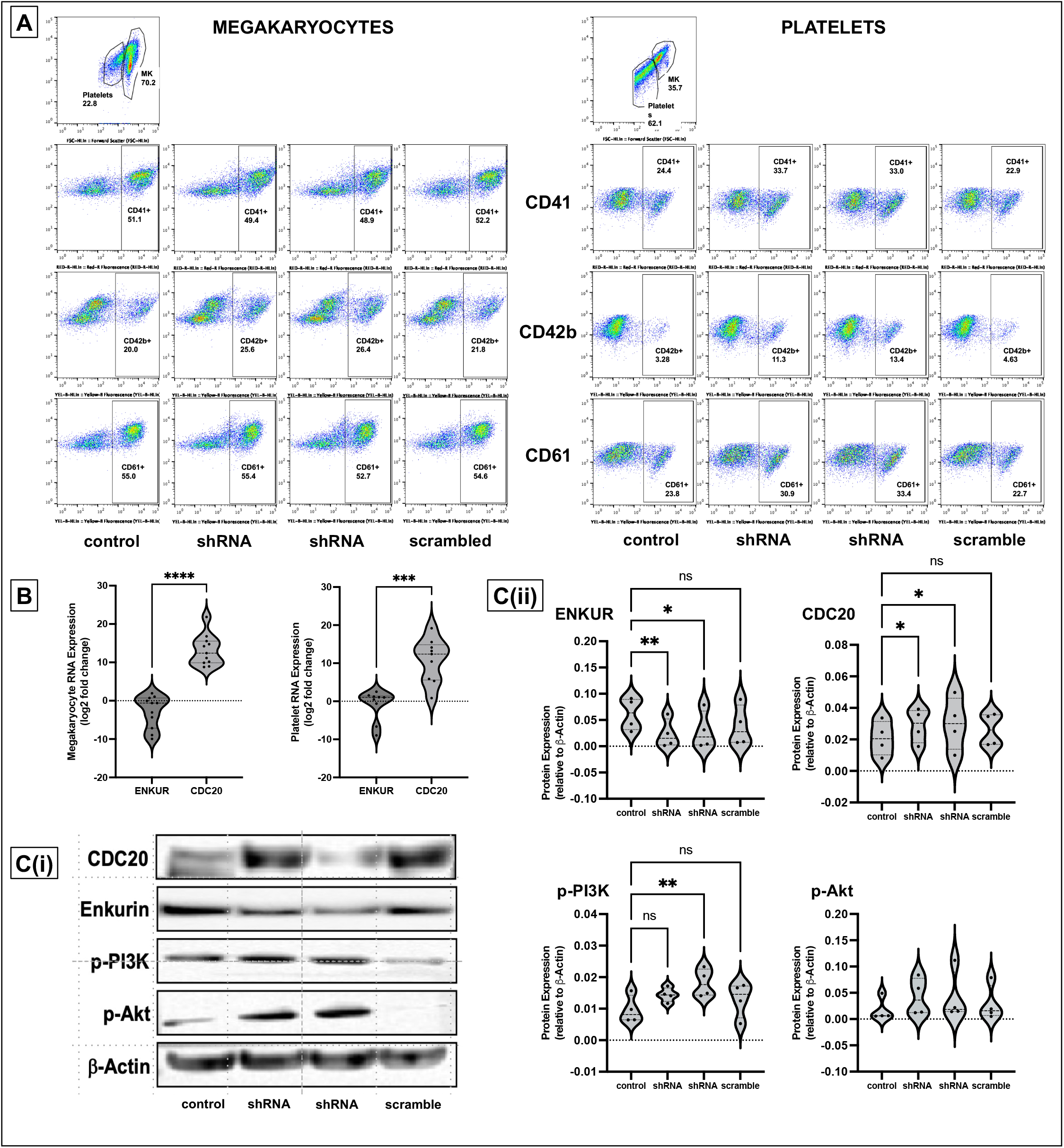
Lentiviral transduction induced silencing of ENKUR gene in CD34^+^ stem cells. (A) Flow cytometric analysis of cell surface markers to confirm the megakaryocytic and platelet fractions after 15 days of culture. (B) RNA expression levels in the CD34^+^ cell-derived megakaryocytes and platelets. (C) (i) Western blot of the CD34^+^ cells derived megakaryocytes (n=4 cord blood and n=4 peripheral blood) and (ii) densitometry for the blots. Densitometry was performed using the IVIS imaging software (****p<0.0001, ***p<0.001, **p<0.01 and *p<0.05 when compared with non-silenced controls).

One classical downstream pathway linked previously to ER stress response^18,19^ in MPNs^52^ and other cancers^53-55^ is the PI3K/Akt signaling cascade. Here with shENKUR, we briefly evaluated possible association with the PI3K/Akt pathway and found that silencing of enkurin correlated with increased expression of phosphorylated PI3K in megakaryocytes (**Figure 4C**).

### Effect of ER stress on CD34^+^-derived megakaryocytes

Considering that the differential and inversely correlated expression of *ENKUR* and *CDC20* extends to the CD34^+^ cell-derived megakaryocytes, and our hypothesis on the role of ER stress in this response, we assessed RNA and protein levels of these markers following treatment of the cells with the sarco/endoplasmic reticulum Ca^2+^-ATPase inhibitor, thapsigargin (125nM). CD34^+^ cells from peripheral blood of healthy donors were differentiated into megakaryocyte and platelet fractions, with thapsigargin introduced at day 7 of a 15-day culture (see Methods and **Figure 5A**; limited cell quantity of the platelet fraction enabled protein analyses in the megakaryocytic fraction alone). Thapsigargin evoked a similar expression profile in megakaryocytes **(Figure 5B-C)** as that of MPN patient bone marrow-derived CD34^+^ cells with downregulated *ENKUR* and upregulated *CDC20* expression at both RNA and protein levels.

**Figure 5:**
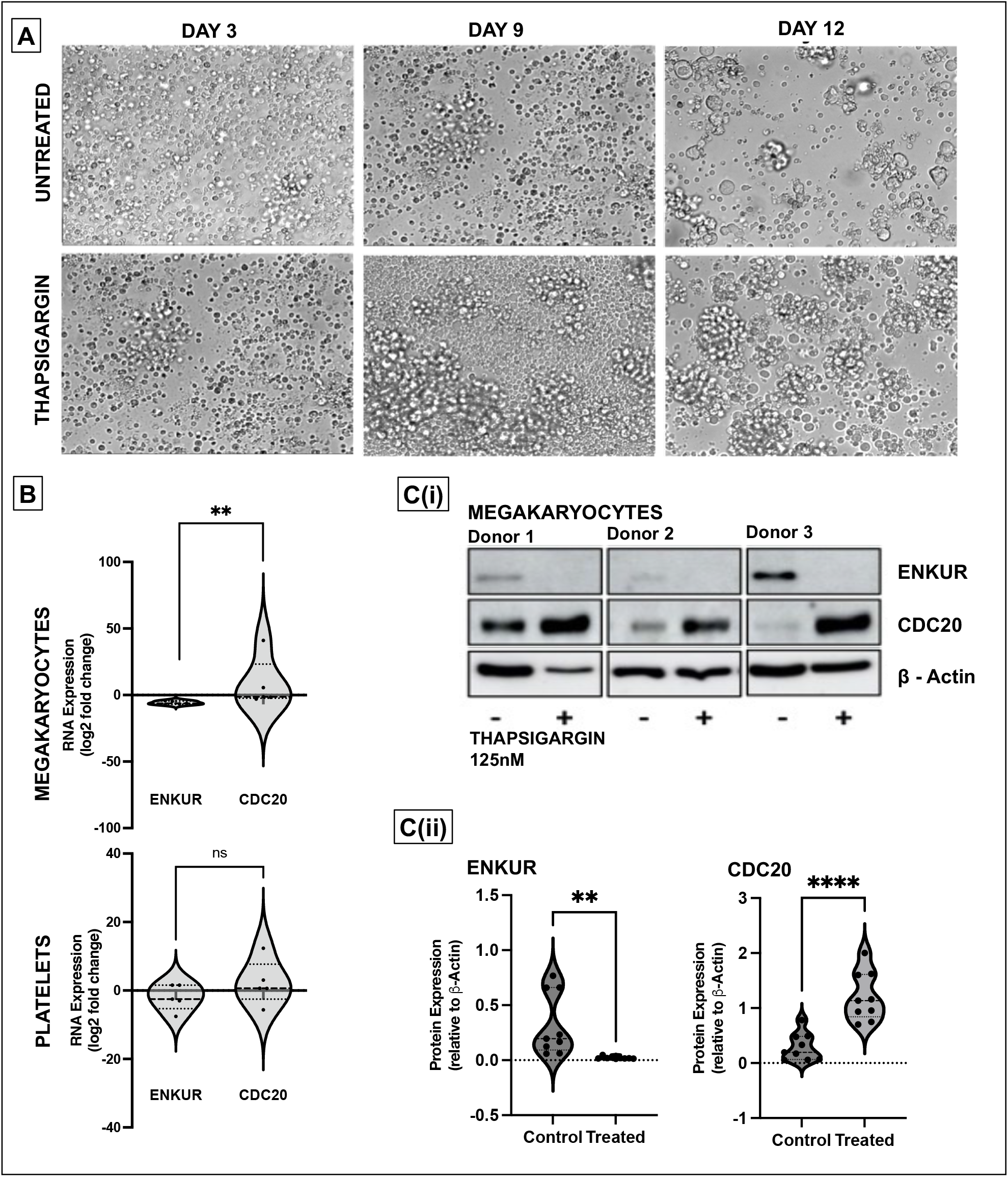
Effect of ER stress on megakaryocytes derived from CD34^+^ cells from healthy donors. (A) Representative microscopic images of the CD34^+^ cells isolated from the peripheral blood of healthy donors over the 15-day culture period at 20X magnification. The ER stressor, thapsigargin (125nM) was added to the culture at day 7 and cultured for another 7 days. (B) RNA expression: cell culture from thapsigargin-treated CD34^+^ derived megakaryocytes (n=7). (C) (i) Western blot of the thapsigargin-treated CD34^+^ cells derived megakaryocytes (n=3 in triplicate) and (ii) densitometry for the blots. Densitometry was performed using the ImageJ software (****p<0.0001,**p<0.01 when compared with untreated controls).

## Discussion

Here, we present the first report, to our knowledge, of enkurin (*ENKUR*) as a potential new peripheral biomarker and therapeutic strategy in myeloproliferative neoplasms. Our prior study^20^ profiling the platelet transcriptome in patients with chronic MPNs (n=120) identified progressive association in expression of several proteostasis-associated genes with advancing disease subtype. Our findings were consistent with other studies demonstrating dysregulated proteostasis as a primary effector of myeloid transformation^18,56,19,57^. The MPN platelet RNA-seq data also confirmed the limited impact^58^ of treatment by JAK2-inhibitor ruxolitinib (RUX) relative to the substantial disease burden in myelofibrosis; and urged the need for development of novel candidate drugs to be used alone or in combination with RUX for the treatment of MPNs.

Four candidate markers from the prior study (*ENKUR, CDC20, CALR, and CREB3L1*) were evaluated in this work based not only on their significant association with pathways related to proteostasis, cell proliferation, ER stress or calcium signaling, but also progressive increase or decrease in differential expression across the MPN chronic vs advanced subtypes (ET/PV to MF versus healthy donors) regardless of patient *JAK2/CALR* mutational status. Using CD34^+^ cells isolated from MPN patient bone marrow as well as healthy donor peripheral and cord blood, we generated *ex vivo* megakaryocyte and platelet fractions and discovered at both RNA and protein levels, an inverse correlation between *ENKUR*, a calmodulin and TRPC channel modulator^26,29^ (not previously associated with MPNs) and *CDC20* (also termed Fizzy^44^), a cell cycle gene and an anaphase promoting complex activator (with known associations with malignancies broadly^43,59^ and select studies in MPNs^60^). Lentiviral-mediated silencing of *ENKUR* was utilized to further confirm the negative association between *ENKUR* and *CDC20;* and indicating a potential interaction via the PI3K-Akt pathway **(Figure 6)**. Our data on the inverse association between enkurin, a membrane calcium influx adaptor protein, and *CDC20*, a crucial cell cycle and proliferation regulator point to a role for dysregulated calcium signaling in MPNs; and the possibility for the ratio of expression between the two to be developed as a potential biomarker of MPN disease.

**Figure 6:**
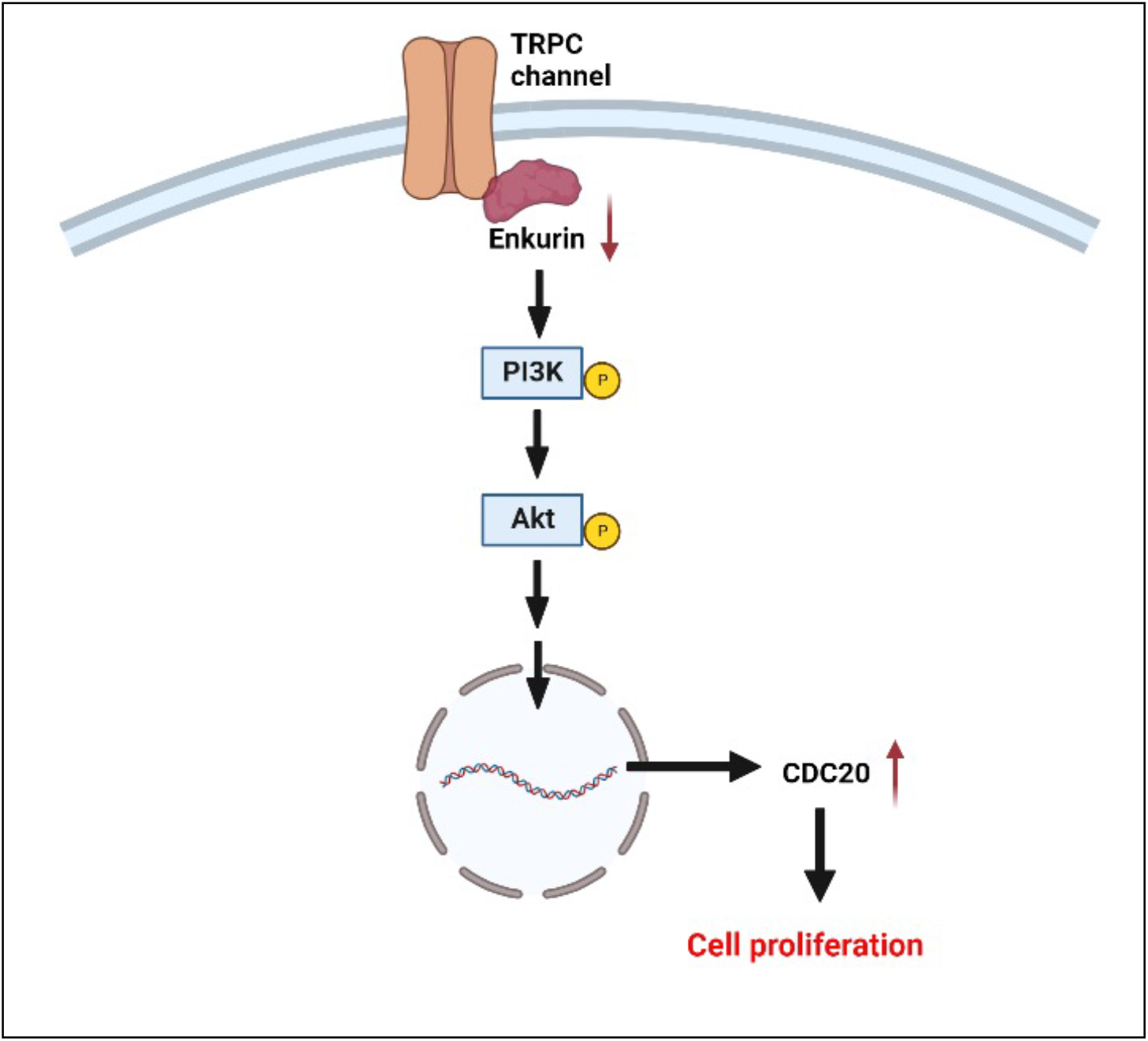
Probable mechanism for the role of *ENKUR* downregulation in cell proliferation. We hypothesize that, under normal conditions, enkurin at sufficient physiological levels is bound to PI3K via the SH3 binding domain, thus regulating its activation^26^. However, in the setting of MPNs, downregulation of enkurin likely contributes to free PI3K and its prolonged activation, further activating Akt and potentially other signaling mechanisms prompting overexpression of cell differentiation cycle genes including CDC20^68^ and cell proliferation.

Next, in evaluating calreticulin (*CALR*), a well-established driver mutation in MPNs^31,61^, as well as *CREB3L1*, an ER/UPR stress response transcription factor, high expression noted previously across MPN patient platelet RNA-seq^20^ was confirmed in patient-derived specimens (platelet, whole-blood, and bone-marrow-CD34^+^-cultured megakaryocytes) but only at the RNA (and not protein) levels. Future investigations applying genome editing in MPN models will be needed to ascertain overlapping mechanisms of somatic alterations in *CALR* and its effect on function in altered calcium signaling and dysregulated ER protein folding^62,63^ in MPNs.

Taken together, we offer two inversely associated MPN markers, *ENKUR* and *CDC20*, whose whole blood, platelet and megakaryocyte RNA and protein expression reflect MPN pathobiology irrespective of patient driver mutational status. These candidate markers also seek to expand our understanding of MPN pathology beyond the classical inflammatory signatures^64-67^.

### Limitations of the study

There are several limitations to our study. First, higher statistical power and longitudinal prospective data will be necessary to further confirm *ENKUR/CDC20* as markers for MPN disease progression. Second, future investigations evaluating these signatures in patient-derived CD34^+^ cells may identify additional functional aspects of bone marrow and MPN pathology. Third, we recognize that our data are not assessing mechanistic signaling of how loss of *ENKUR* expression or impaired calcium signaling contributes to myeloproliferation; and will need to be specifically interrogated in MPN murine and other models. Follow-on studies evaluating a potential role of the *PI3K/Akt* pathway, in particular the effect of pharmaceutical inhibitors already in use in MF will add significant value to the current study. Studies evaluating platelet function under impaired enkurin expression and calcium modulation are promising immediate future directions. And above all, it will be important to assess how enkurin downregulation might be more broadly relevant to other hematological cancers, such as myelodysplastic syndromes and acute myeloid leukemia, or potentially unique to MPNs alone.

## Acknowledgements

This work was funded by the MPN Research Foundation and the US National Institutes of Health grants 1K08HG010061-01A1 and 3UL1TR001085-04S1 (research re-entry award) to A.K. Other support included 1S10OD018220 and 1S10OD021763 shared instrumentation grants to the Stanford Functional Genomics Facility and the Stanford Research Computing Center, and the Charles and Ann Johnson Foundation to J.G. All authors thank the patients at the Stanford Cancer Center and the healthy donors at the Stanford Blood Center for their generous participation in this research, and the Canary Center at Stanford for Cancer Early Detection for access to flow cytometry and microscopy. Authors also thank Dr. Diwash Jangam for his assistance with the initial Paxgene PCR setup.

## Author Contributions

AK conceived of the study and the team, secured funding, and wrote the manuscript. LF and JG provided samples and clinical annotation and reviewed the clinical data. AK and RP coordinated and oversaw sample acquisition and processing. SS, YL, JW, and KM performed the experiments and interpreted the analyses with AK, RP, HM, JG, and JZ. All authors critically reviewed and edited the manuscript. ^#^SS and YL contributed equally. ^#^JG and JZ also contributed equally. All authors approved the final manuscript.

## Supplementary Figure legends

**Figure S1:**
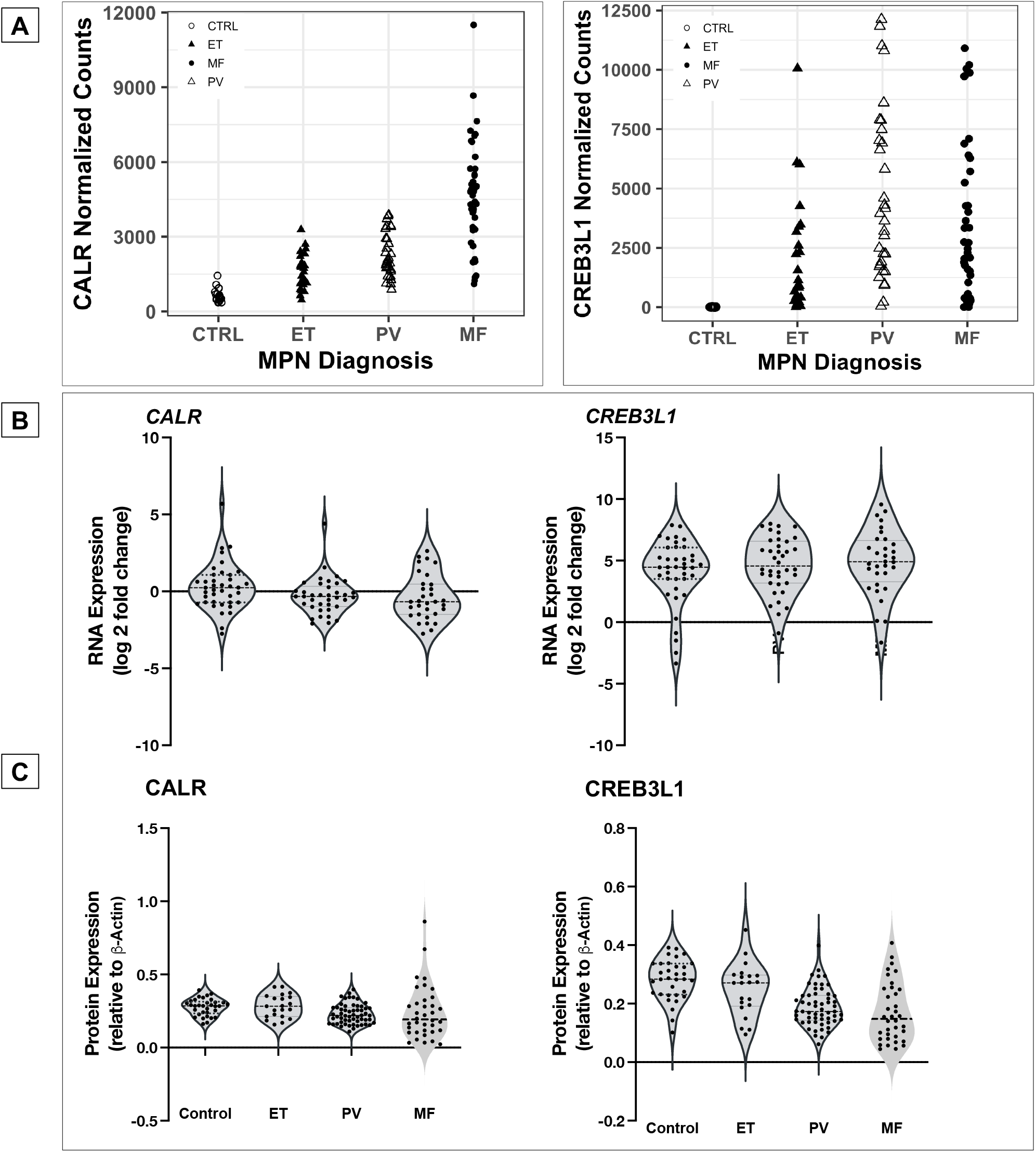
RNA expression in platelets and whole blood of MPN patients and protein expression in platelets. (A) MPN Platelet RNA-seq Normalized Expression Data: MPN patients (n=99: ET n=24; PV n=33; MF n=42) vs healthy donors (n=21) (B) whole blood RNA: MPN patients (n=109: ET n=39; PV n=38; MF n=32) vs healthy donors (n=10) (C) platelet protein: peripheral blood platelets were isolated from MPN patients (n=38: MF n=12; PV n=19; ET n=7) and healthy donors (n=11).

**Figure S2:**
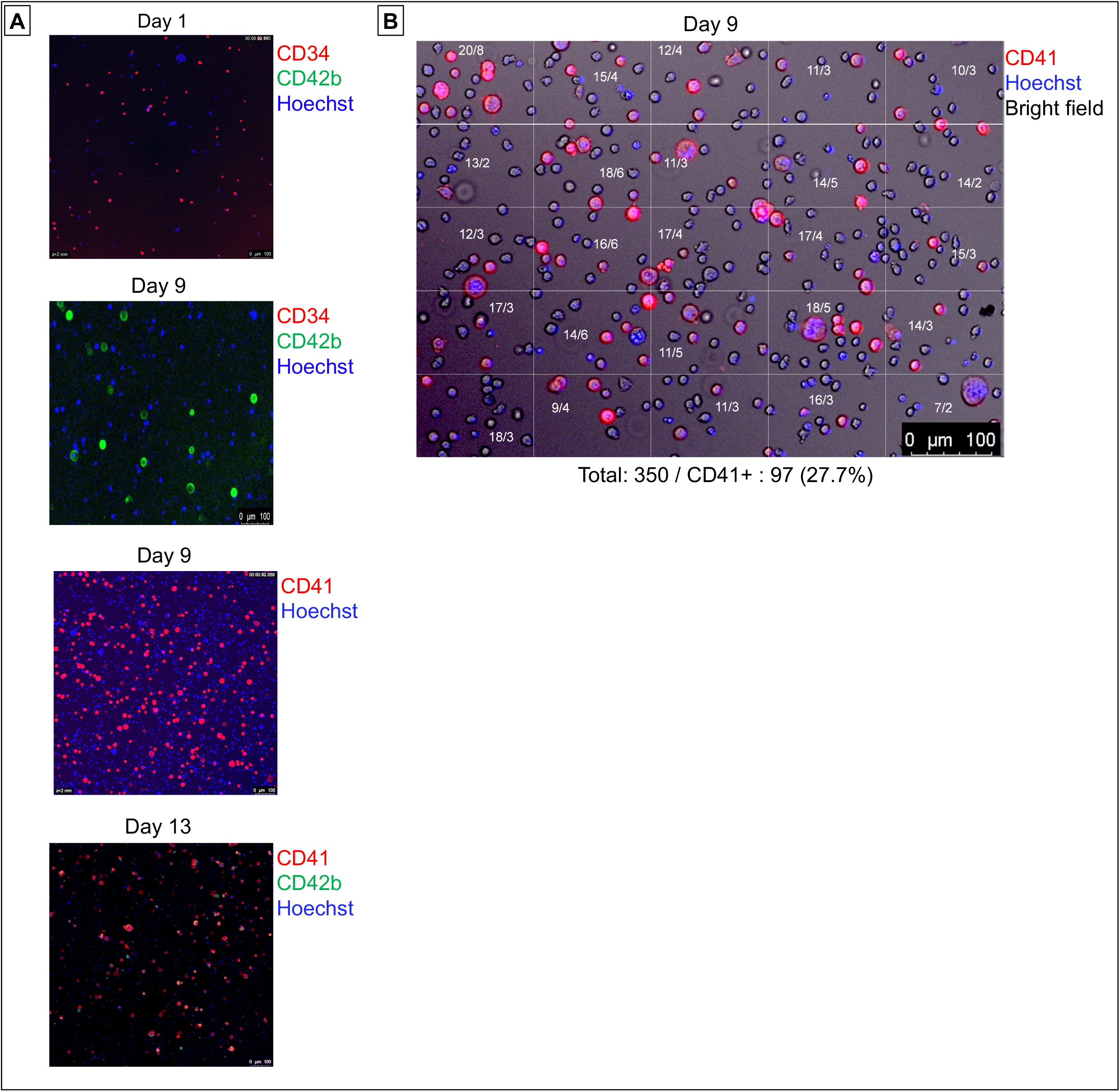
Culturing of CD34^+^ cells from the bone marrow of MPN patients and RNA expression. (A) Confocal microscopy: Microscopic images of the CD34^+^ cells isolated from the bone marrow of MPN patients over the 15-day culture period at 20X magnification. The cells were treated with cell surface markers for CD34^+^ cells (CD34), Megakaryocytes (CD41, CD42b, and CD61) (B) Quantification of CD41^+^ megakaryocytes from CD34^+^ cells isolated from the bone marrow of MPN patients over the 15-day culture period (C) Megakaryocytic RNA: cell culture from fresh MPN patient bone marrow (n=8) derived CD34^+^ cells then differentiated over 15 days into megakaryocytes. All RNA expression is normalized to GAPDH and expressed as log2 fold change.

**Figure S3:**
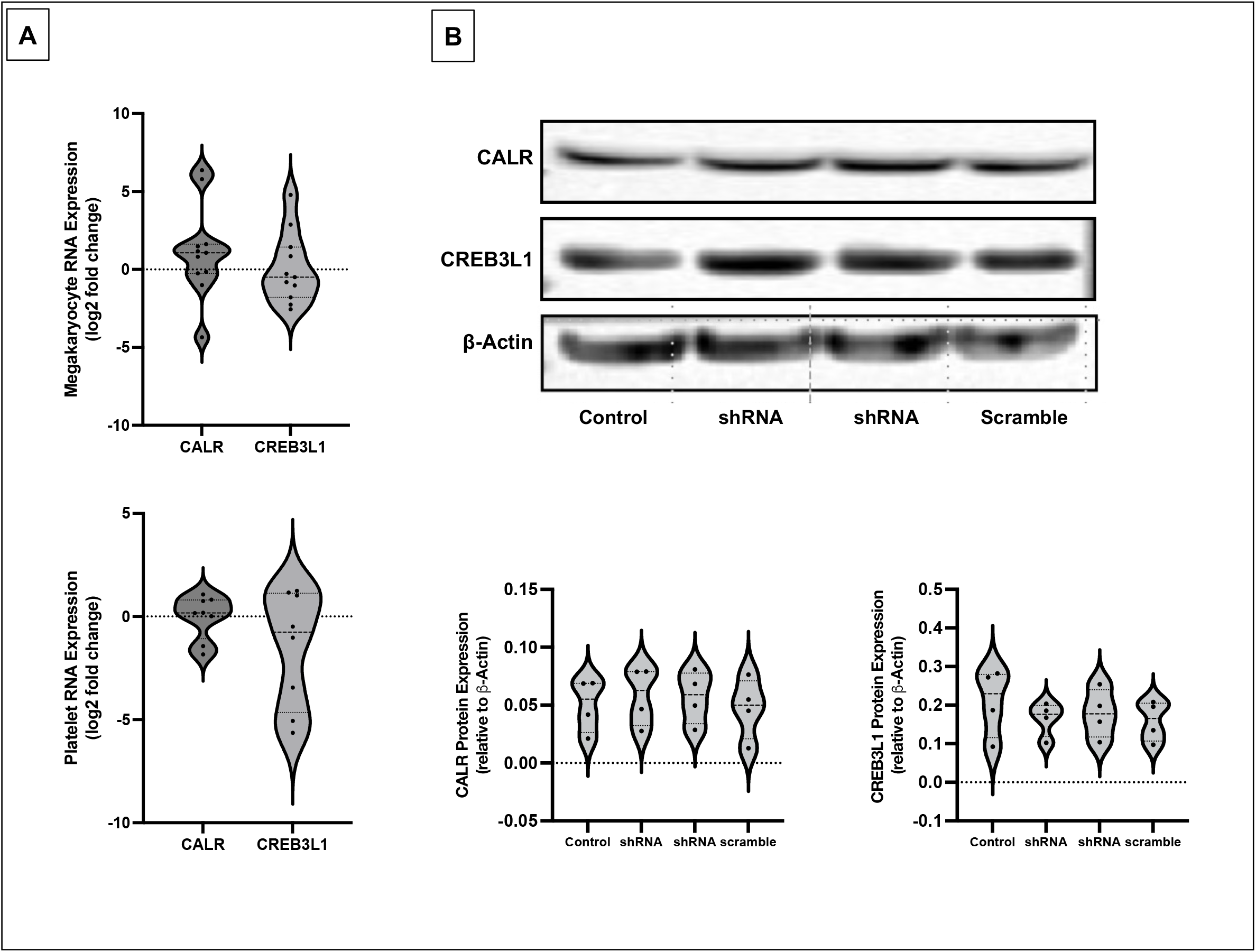
Effect of Lentiviral transduction induced silencing of ENKUR gene on CALR and CREB3L1 gene expression in CD34^+^ stem cells. (A) RNA expression levels in the CD34^+^ cell-derived megakaryocytes and platelets. Log2 fold change was plotted by normalizing with the control (B) (i) Western blot of the CD34^+^ cell-derived megakaryocytes (n=4 cord blood and n=4 peripheral blood) and (ii) densitometry for the blots. Densitometry was performed using IVIS imaging software.

